# Towards the human cellular microRNAome

**DOI:** 10.1101/120394

**Authors:** Matthew N. McCall, Min-Sik Kim, Mohammed Adil, Arun H. Patil, Yin Lu, Christopher J. Mitchell, Pamela Leal-Rojas, Jinchong Xu, Manoj Kumar, Valina L. Dawson, Ted M. Dawson, Alexander S. Baras, Avi Z. Rosenberg, Dan E. Arking, Kathleen H. Burns, Akhilesh Pandey, Marc K. Halushka

## Abstract

microRNAs are short RNAs that serve as master regulators of gene expression and are essential components of normal development as well as modulators of disease. MicroRNAs generally act cell autonomously and thus their localization to specific cell types is needed to guide our understanding of microRNA activity. Current tissue-level data has caused considerable confusion and comprehensive cell-level data does not yet exist. Here we establish the landscape of human cell-specific microRNA expression. This project evaluated 8 billion small RNA-seq reads from 46 primary cell types, 42 cancer or immortalized cell lines, and 26 tissues. It identified both specific and ubiquitous patterns of expression that strongly correlate with adjacent super-enhancer activity. Analysis of unaligned RNA reads uncovered 207 unknown minor strand (passenger) microRNAs of known microRNA loci and 2,632 novel putative microRNA loci. Although cancer cell lines generally recapitulated the expression patterns of matched primary cells, their isomiR sequence families exhibited increased disorder suggesting Drosha and Dicer-dependent microRNA processing variability. Cell-specific patterns of microRNA expression were used to deconvolute variable cellular composition of adipose tissue samples highlighting one use of this cell-specific microRNA expression data. Characterization of cellular microRNA expression across a wide variety of cell types provides a new understanding of this critical regulatory RNA species.

## Introduction

MicroRNAs are an established class of small regulatory RNAs that, within the RISC complex, bind mRNAs and repress protein production (Valencia-Sanchez et al. 2006). In this role, they control essential cell processes in health and disease (Ambros 2004; Mendell and Olson 2012). Despite all that is known about microRNA processing and function, the cellular localization of microRNAs is still widely underappreciated. An understanding of which cells express which microRNAs is useful as we move towards microRNA therapeutics (Janssen et al.2013) and microRNA biomarkers (Mitchell et al. 2008). Knowing a microRNA’s full localization pattern will maximize efficacy and minimize off-target effects of therapeutics and will better rationalize candidate biomarkers (Haider et al. 2014).

MicroRNA expression has been predominately characterized in tissues, with no comprehensive cellular studies. Initial tissue studies sequenced individual clones or used array methods providing low-depth coverage of abundant microRNAs (Lagos-Quintana et al. 2002; Barad et al. 2004; Liu et al. 2004; Baskerville and Bartel 2005). The most thorough of these microRNA localization projects performed small RNA library sequencing (RNA-seq) on over 250 libraries from 26 organ systems. But this nascent effort sequenced only ∼1,200 reads per library (Landgraf et al. 2007). While providing valuable insights into the relationship of microRNA expression and disease (Lu et al. 2005), these and subsequent studies (Cheng et al. 2015; Ludwig et al. 2016) have not unraveled cellular microRNA expression. Because all tissues are composed of multiple, unique cell types, it is essential to understand from which cell the microRNA signal is obtained. Additionally, the anonymity of microRNA nomenclature, with sequential numerical naming has not allowed an intrinsic understanding of what microRNAs are ubiquitous and which have cell-restricted patterns of expression (Witwer and Halushka 2016). This determination is fundamental to understanding the proper biologic and regulatory roles of microRNAs.

Small RNA-seq has become a robust method to fully characterize known microRNA, capture complete isomiR families, and identify novel microRNAs. IsomiRs are related sequences with mostly 5’ and 3’ nucleotide modifications that collectively make up the totality of a given microRNA (Neilsen et al. 2012). The microRNA community has been forthright in depositing RNA-seq data into central public repositories. As a result, there is a significant amount of data that can be collectively analyzed. We combined new sequencing of 39 primary cell lines or isolated cells with hundreds of publicly available primary cell and immortalized/cancer cell line datasets, with all microRNA assignment performed by a single robust and high-throughput microRNA alignment method (Baras et al. 2015) to establish the most complete characterization of the human cellular microRNAome including novel microRNA discovery and isomiR diversity. We additionally analyzed whole tissue microRNA data to understand the extent to which cells obtained from *ex vivo* cultures could recapitulate a tissue signal and compared matched primary and cancer/immortalized cells to determine the extent of similarity in their expression patterns.

## Results

### Generation of a cellular microRNAome

Towards cataloguing a high-quality complete cellular microRNAome, we generated new small RNA-seq data from 39 primary cells obtained by culture, flow cytometry or centrifugation. We augmented this with Sequence Read Archive (SRA) small RNA-seq read data from 496 samples with > 1 million microRNA reads. These were primary cell cultures, immortalized/cancer cell lines or normal tissues (Fig. 1). All samples were processed through miRge (Baras et al. 2015). miRge uses modified microRNA libraries and multiple Bowtie steps for optimal alignments on multiplexed runs. (Fig. 2A, Table 1). Overall, 2,319 of 2,546 known microRNAs (miRBase v21) had a minimum expression of 1 read per million microRNA reads (RPM) in at least one sample (Supplemental Table S1).

**Figure 1.**
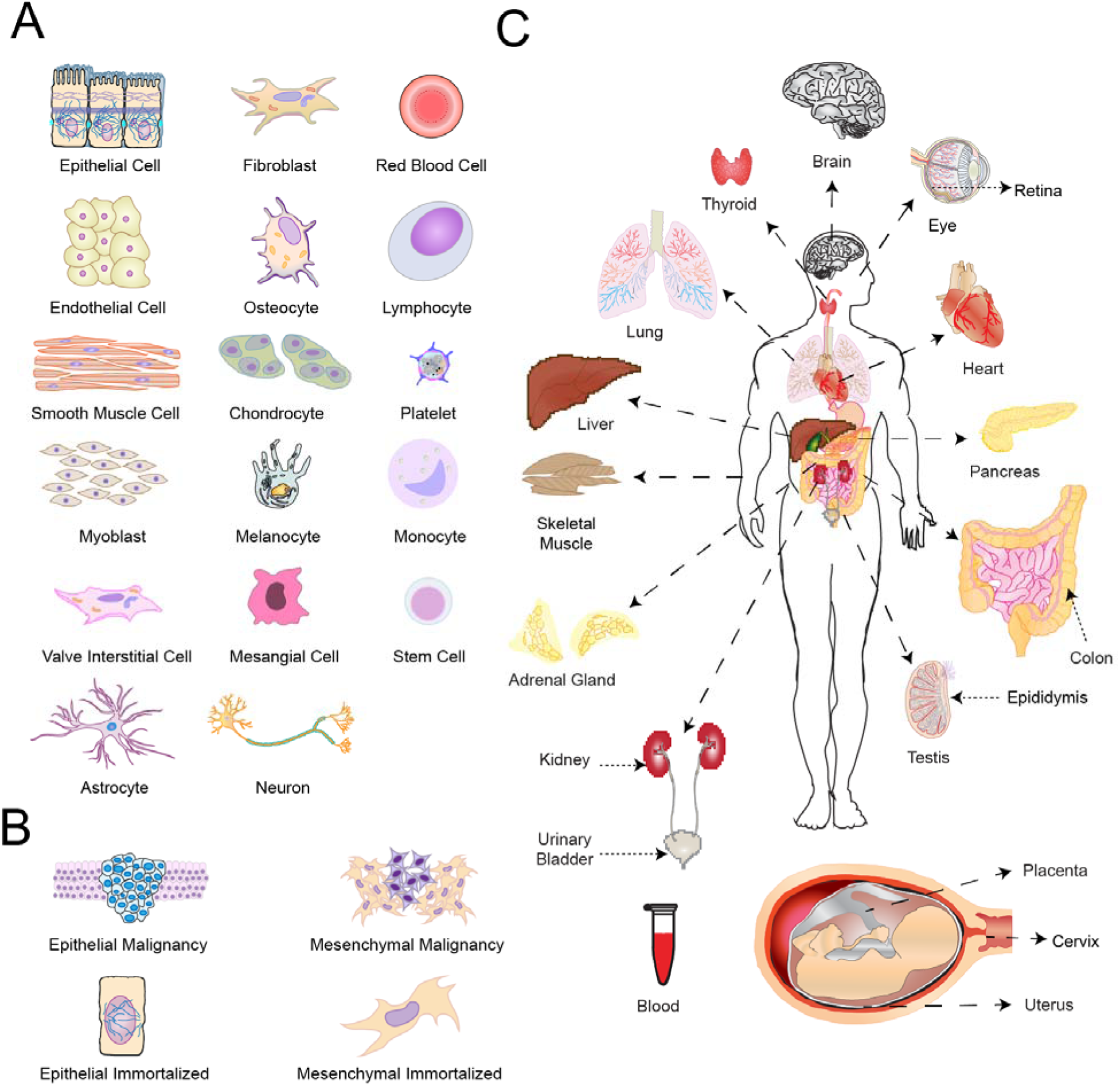
A generalized overview of the 530 cells and tissues included in this study. (A) Representation of 46 main cell types (B) Representation of 42 cancer or immortalized cell lines (C) Representation of 26 tissues/organ types.

**Figure 2.**
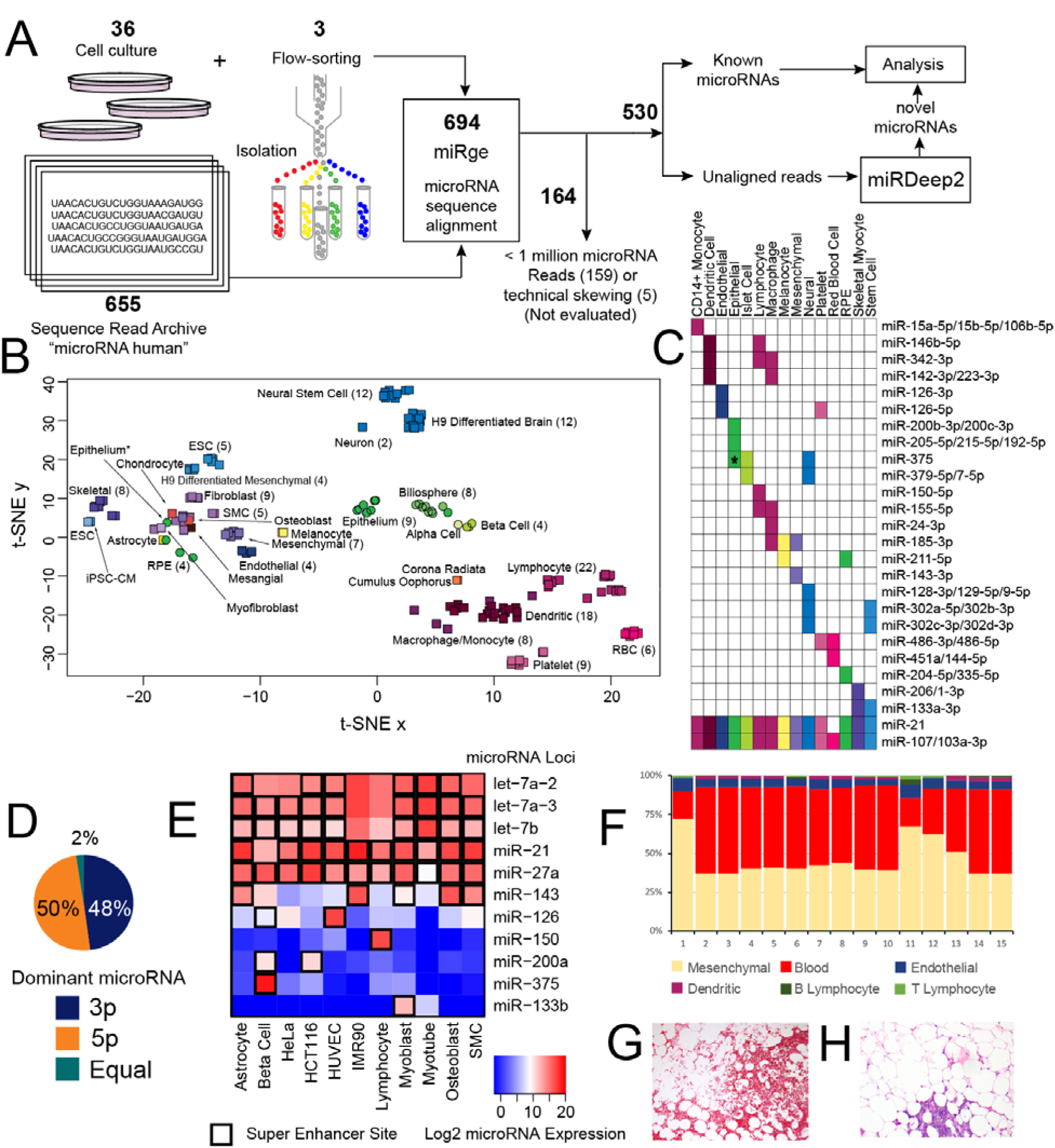
Method and primary analysis of the cellular distribution of microRNAs. (A) 694 total samples were processed through miRge yielding 530 samples available for analysis and novel microRNA detection through miRDeep2. (B) t-SNE distribution of 161 primary cells showing 4 main clusters (hematologic, epithelial, mesenchymal and neural/stem cell) and subclustering by cell type. Cell types are color-coded and round symbols indicate epithelial cells. * indicates an intestinal epithelial cell that was either contaminated or underwent mesenchymal transformation. (C) A selection of microRNAs that have unique expression to certain primary cell types. * indicates specificity for flow-sorted colonic epithelial (likely goblet) cells. (D) Across 334 microRNAs with >1,000 RPM, both strands of a hairpin microRNA give rise to the dominant microRNA in fairly equal measures. (E) The presence of nearby super-enhancers strongly correlates with high microRNA expression. (F) The individual cellular microRNA patterns can be used to deconvolute the cellular composition of tissue. (G) A representative hematoxylin & eosin (H&E) section of adipose with significant red blood cells (lower part of the panel) as an example of heterogenous elements that can contribute microRNA expression (10x original magnification). (H) A H&E representative section of adipose with a small cluster of lymphocytes (lower part of the panel) that may be randomly sampled modulating the tissue signal (10x original magnification).

**Table 1.**
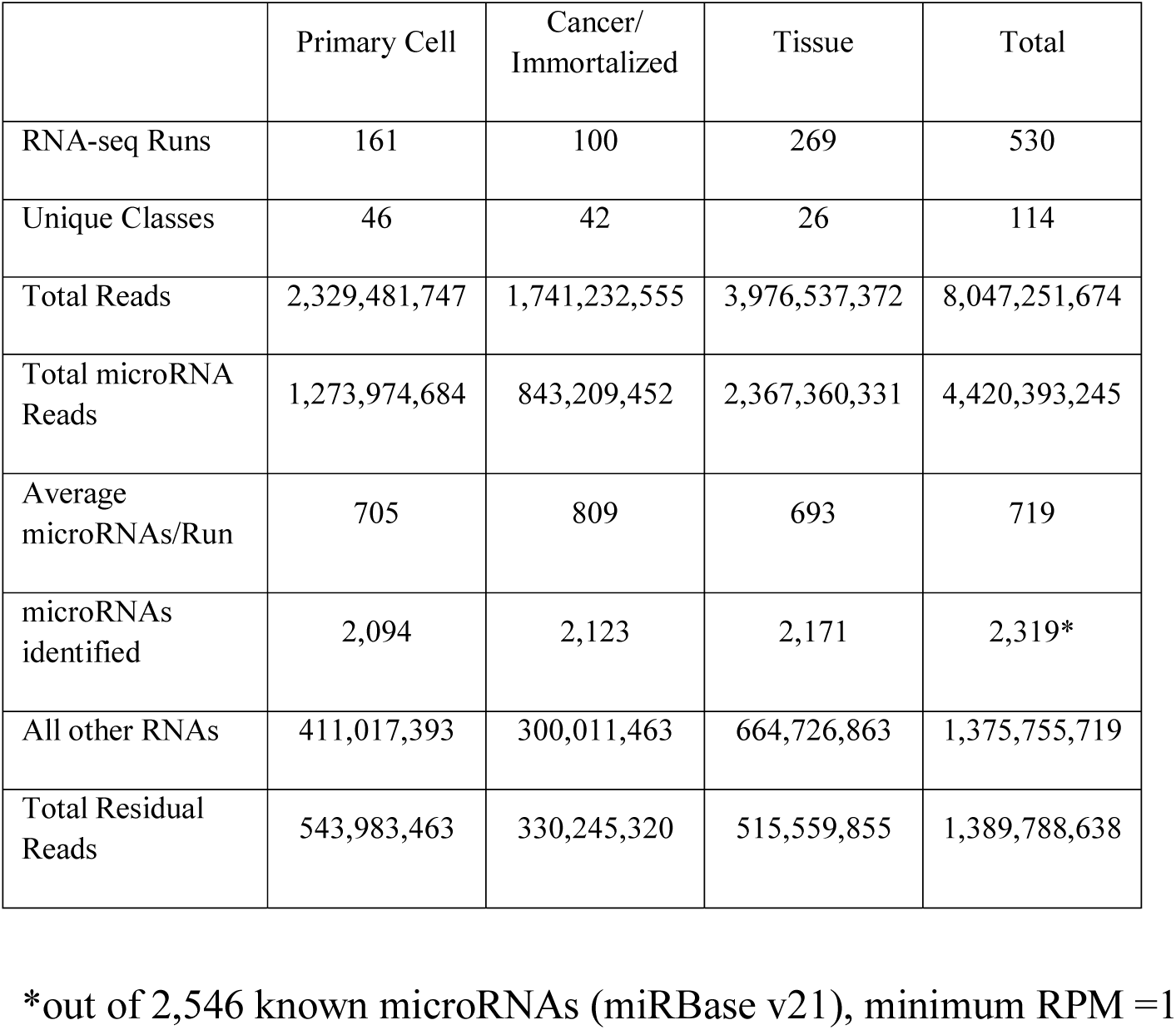
Overall Sequencing Data.

|  | Primary Cell | Cancer/<br>Immortalized | Tissue | Total |
| --- | --- | --- | --- | --- |
| RNA-seq Runs | 161 | 100 | 269 | 530 |
| Unique Classes | 46 | 42 | 26 | 114 |
| Total Reads | 2,329,481,747 | 1,741,232,555 | 3,976,537,372 | 8,047,251,674 |
| Total microRNA Reads | 1,273,974,684 | 843,209,452 | 2,367,360,331 | 4,420,393,245 |
| Average microRNAs/Run | 705 | 809 | 693 | 719 |
| microRNAs identified | 2,094 | 2,123 | 2,171 | 2,319* |
| All other RNAs | 411,017,393 | 300,011,463 | 664,726,863 | 1,375,755,719 |
| Total Residual Reads | 543,983,463 | 330,245,320 | 515,559,855 | 1,389,788,638 |

\*out of 2,546 known microRNAs (miRBase v21), minimum RPM =1

The 161 primary cell RNA-seq data sets encompassed 46 main cell types, many from multiple anatomic locations (Fig. 1A, Supplemental Table S2). There were 100 cancer cell or immortalized cell line RNA-seq data sets from 42 separate cell lines (19 general cancer types) (Fig. 1B, Supplemental Table S3). The 269 small RNA-seq data sets from 26 normal tissues/organs aided in the normalization methods employed due to organ coverage from multiple separate studies (Fig. 1C, Supplemental Table S4). As much of this primary data was derived from different laboratories using different protocols, significant attention was given to potential confounding and batch-effects.

We utilized the DEXUS algorithm (Klambauer et al. 2013), to identify discrete expression states for each microRNA. The resulting cell-type specific patterns of discretized microRNA expression across the 161 primary cell types (Supplemental Fig. S1) are inherently robust to batch effects. This method clustered cell types into hematologic, neural/embryonic stem cell (ESC), epithelial and mesenchymal groups, identifying general patterns of microRNA expression. T-distributed Stochastic Neighbor Embedding (t-SNE) clustering was then compared between uncorrected primary cell RPM data and primary cell data that underwent Remove Unwanted Variation (RUV) normalization for 5 variables using the most abundant microRNAs (Risso et al. 2014). The use of RUV improved clustering of similar cell types from different experiments (Supplemental Fig. S2).

t-SNE was then performed separately for primary cells, cancer/immortalized cells and tissues using RUV normalization (Fig. 2B, Supplemental Fig. S3). Akin to the DEXUS results, amongst primary cell types, microRNA expression patterns generated four major groups: hematologic, mesenchymal, neural/ESC and epithelial. Strong clustering by biological group was observed for all samples, overcoming most technical concerns.

### Diverse microRNAs expression patterns

We assessed common, potentially functional microRNAs (Mullokandov et al. 2012) by their frequency of expression across the different normal cell classes (Supplemental Table S5). There were 320 microRNAs that had an RPM ≥ 1,000 in any of the 46 normal cell types. Of these, 94 (29%) were present in only a single class of cells (Fig. 2C). Most of these are well-known associations (ex. miR-144-3p and red blood cells or miR-1-3p with skeletal myocytes) that highlight the non-ubiquitous nature of microRNA expression (Haider et al. 2014). Six microRNAs were present in all 46 cell types at this RPM threshold (miR-107, miR-103a-3p, miR-103b, miR-191-5p, miR-21-5p, and miR-92a-3p) and an additional 9 microRNAs were present in all cells at a lower threshold of 100 RPM: miR-16-5p, miR-25-3p, miR-26a-5p, miR-26b-5p, miR-30d-5p, miR-101-3p, miR-128-3p, miR-140-3p, and miR-181a-5p (Supplemental Table S6). Among tissues containing a mixture of cell types, 377 microRNAs were present at an RPM ≥ 1,000. Seven of these microRNAs (let-7a-5p, let-7c-5p, let-7b-5p, let-7f-5p, let-7g-5p, miR-26a-5p, miR-30d-5p) were found in all tissues. Some well-known cell-specific microRNAs appear to be ubiquitous amongst tissue, but merely reflect the presence of a certain cell type across tissues. miR-451a was abundantly present in 20 of 26 tissues but is from only one cell class (red blood cells). Likewise, miR-126-3p, abundant only in endothelial cells, was present in 21 tissues and miR-150-5p, abundant in lymphocytes, was present in 11 tissues. We then determined microRNA abundance from the 5p or 3p arm and found the guide/“driver” (more abundant, thermodynamically-stable) microRNA to be equally from either arm of the hairpin microRNA suggesting no strand bias in microRNA selection (Fig. 2D).

Super-enhancers are dense genomic regions of transcription factor binding sites that have a multiplicative effect on increasing adjacent gene expression (Whyte et al. 2013). We examined the association between microRNA expression levels and the presence of a super-enhancer within 40kb of the microRNA loci in 11 primary cells and cancer cell lines, for which we had matching data. MicroRNA expression in the presence of a super-enhancer was significantly increased compared to microRNA expression at sites not adjacent to a super-enhancer (Wilcoxon rank sum test p<2.2^-16^). This association was observed with and without batch effect adjustment and was generally consistent across samples. Importantly, cell-type restricted microRNAs showed active super-enhancer activity matching those specific cells (Fig. 2E). These data further support the cell-type restricted microRNAs seen above and indicate that analyses of global tissue microRNA expression require an understanding of the source of each microRNA of interest, lest misinterpretations of the data result in spurious disease associations (Kent et al. 2014).

Taking advantage of cell-specific microRNAs, we determined the feasibility of using cellular microRNA expression data to deconvolute an overall tissue signal to discern the individual components. We investigated 15 white adipose samples from participants in the METSIM study (Civelek et al. 2013). We used CIBERSORT (Newman et al. 2015) to determine the composition of each tissue based on grouped microRNA signatures for mesenchymal cells, endothelial cells, blood, B/T lymphocytes and dendritic cells (Fig. 2F). Surprisingly, the red blood cell component (blood) was a major and variable (18-56%) part of each tissue suggesting inconsistent and inadequate sample washing prior to RNA isolation (Fig. 2G). Lymphocytes were also variable (0-5%) between samples (Fig. 2H), while endothelial cells were generally more consistent (4-9%). Altogether, these data demonstrate the importance and feasibility of solving for the cellular content of tissues to better understand the composition of the analyzed tissue.

### Variable expression in cancer cell lines

Immortalized and cancer-derived cell lines are frequent surrogates for primary cells in understanding biologic pathways. However, the extent of differences in microRNA expression between these cell lines and primary cells is unknown. We analyzed fibroblasts and T lymphocytes, the only two cell types in which there exists sufficient numbers of primary and immortalized/cancer cell types to determine the extent of their microRNA similarities. An analysis of 12 primary and 3 immortalized fibroblast cell line microRNA signatures identified overall strong correlation (RUV corrected, log2 normalized, pairwise R >0.80-0.99) with the immortalized lines being slightly less correlated (Supplemental Fig. S4A). A global comparison of microRNA differences identified only miR-1304-3p to be significantly different among fibroblasts (Supplemental Fig. S4B). Eight primary T lymphocyte samples and 14 T lymphocyte malignancy samples also revealed moderate to strong correlation, but with separate primary and cancer-derived cell clustering (Supplemental Fig. S4C). There were more microRNAs that differed between primary and cancer-derived cells (Supplemental Fig. S4D) including miR-150-5p which was 6 log2-fold higher in the primary T cells, as has been reported (He et al. 2014). miR-9-3p, 6 log2-fold higher in the cancer-derived cells, has been previously reported as elevated in Hodgkins lymphoma (Leucci et al. 2012) but not in these three cell types. Other markedly different, well-studied, microRNAs include miR-363-3p, miR-146a-5p and miR-146b-5p and miR-486-3p. We also ascertained how consistent the microRNA expression pattern of cancer cell lines would be after years of divergent growth in separate laboratories. A comparison of HeLa cells obtained from 5 sources had a range of expression correlation between 0.35-0.75, while fibroblasts from 3 separate batches, but obtained from different organ systems, had a correlation between 0.75-0.9 (Supplemental Fig. S5). These analyses suggest some key differences between immortalized/cancer cell lines and primary cells and highlight NIH concerns about the rigor and reproducibility of widely used cancer cell lines.

### Novel human microRNAs

We investigated 1.2 billion reads, unmapped by miRge, from 474 samples for putative novel microRNAs in miRDeep2 (Friedlander et al. 2012). miRDeep2 identified, and we assigned names (JHU_ID_xxx), to 25,218 putative “driver” (thermodynamically stable) and “passenger” (thermodynamically unstable and degraded) microRNAs from 21,338 loci with the majority (18,480/65%) being from individual samples and frequently (5,662/22%) identified from only a single read (Supplemental Table S7A). A small percent (394, 0.7%) were identified in more than 50 samples and 207 were the unassigned “passenger” 5p or 3p microRNA from a known microRNA locus (Fig. 1D, Supplemental Table S8). Because most of the rare novel microRNAs likely represent false positives and/or nonfunctional transcripts,(Mullokandov et al. 2012) we selected only the 2,724 loci containing 4,064 mature microRNAs that had ≥50 combined dominant microRNA reads and to consider further (Supplemental Table S9).

Of these 2,724 microRNA loci, 2,632 were completely novel loci and 92 were either the “passenger” strand of known human microRNAs (n= 77) or orthologous (n = 15; Supplemental Table S10) to a different species’ microRNA (primarily primate). We checked the distribution of z-scores for these (and a subset of 21,338 loci) using novoMiRank and found that, the z-scores were right shifted, indicating less similarity to known microRNAs based on 24 features of comparison (Supplemental Fig. S7B,C) (Backes et al. 2016). For these 2,632 putative novel microRNAs, 1,347 (51%) were located within a gene locus and 159 of these microRNAs were also adjacent to a known microRNA. These 2,724 putative microRNA loci were compared to 3,707 novel microRNAs recently reported (Londin et al. 2015) and 21,908 putative novel microRNAs obtained from 105 Argonaute Clip-seq data sets (Ago) run through miRDeep2 and were found to frequently match these two data sets (667/24% to the Londin microRNAs; 1,489/55% of the Ago microRNAs) with 561 microRNAs shared between all three groups (Fig. 3A). We also investigated if these putative novel microRNAs overlapped repeat elements based on the RepeatMasker track of the UCSC Genome browser and indeed 900 (33%) did. Although this would suggest they were less likely to be true microRNAs, 46% and 20% of these putative novel microRNAs were also detected in the Ago and Londin data sets. As well, removing these 900 putative novel microRNAs did not change the average NovoMiRank z-score (1.03 vs 1.03). The median number of reads per “driver” microRNA in this limited sample was 166 (range 50 - 2,662,374).

**Figure 3.**
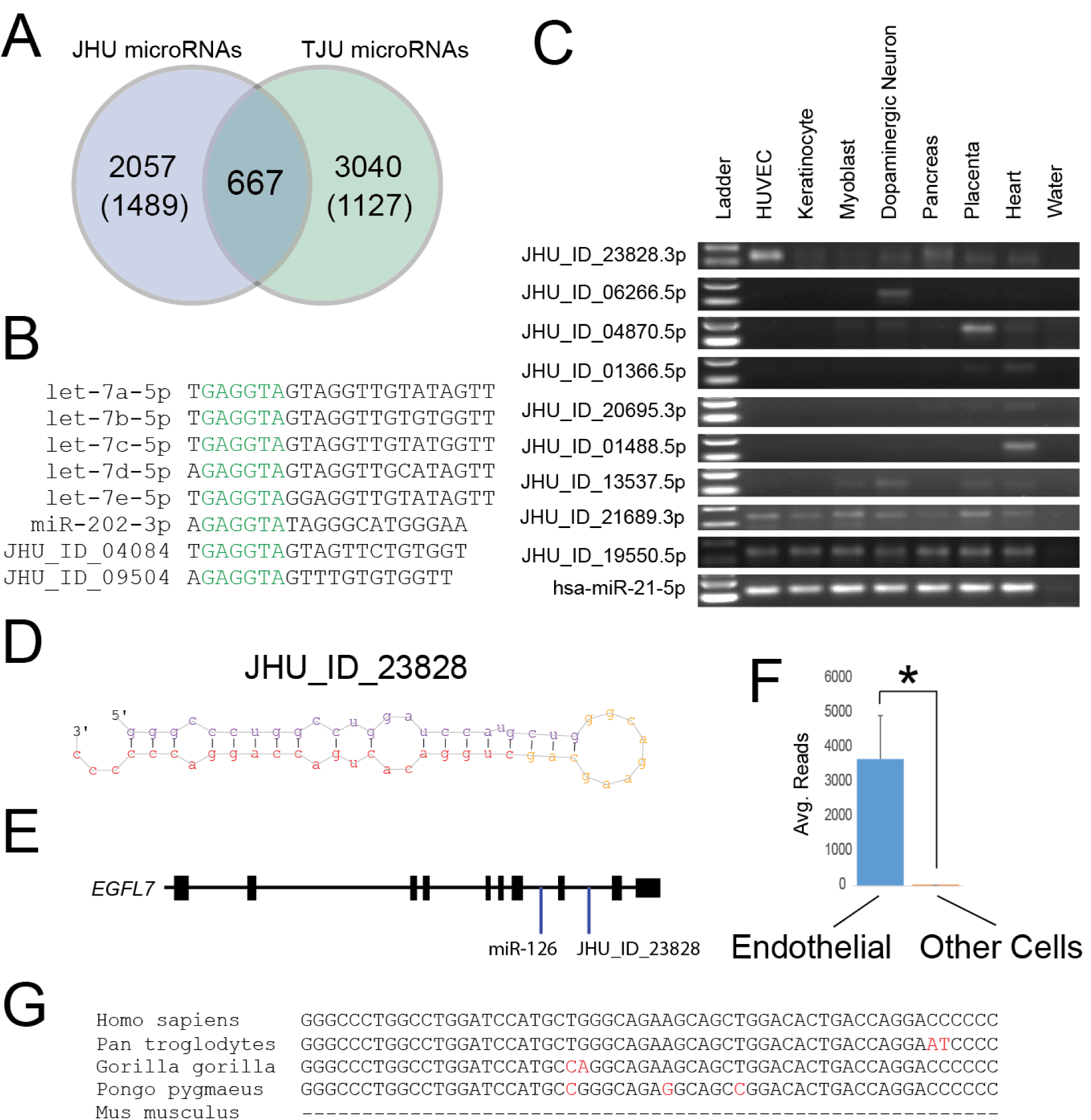
Interesting novel human microRNAs still need to be characterized. (A) A Venn Diagram shows the overlap between putative microRNAs from this publication (JHU) and novel microRNAs reported in Londin *et al* (TJU). Numbers in parentheses represent microRNAs that also overlapped Ago-bound putative microRNAs. (B) 8 representative microRNAs that share the GAGGUA motif. While the let-7 family is ubiquitously expressed, miR-202-3p and JHU_ID_04084 are highly expressed in testis. (C) 9 novel microRNAs were amplified that were predicted to be either cell-specific or ubiquitous. Most of these were lowly expressed. miR-21-5p was used as a control. (D) The predicted hairpin structure of novel microRNA JHU_ID_23828 is shown. (E) JHU_ID_23828 is located in the *EGFL7* gene locus and shares a pre-miRNA with miR-126, an endothelial cell-enriched microRNA. (F) From the same sequencing batch, the average number of reads for JHU_ID_23828 among 4 endothelial cell types was 3,673 and 8 among 29 non-endothelial cell types. (*p=0.001, Mann-Whitney U test) (G) JHU_ID_23823 is present among primate species but is absent in lower mammals including *mus musculus*.

Total reads per microRNA did not correlate with the number of samples within which the microRNA was detected (R^2^<0.01) (Supplemental Fig. S6). The median number of samples sharing a given microRNA loci was 11 (range 1 – 376) (Supplemental Fig. S7). We then investigated similarities by seed regions (bases 2-7 of a microRNA). There were 2,003 novel microRNAs that shared 803 seed regions with known microRNAs. This included 5 novel microRNAs with a “GAGGUA” seed that matched the common seed of 12 let-7 members and 2 novel microRNAs that shared the common “AAGUGC” seed of 15 members of the miR-302/miR-519/miR-520 family (Fig. 3B, Supplemental Table S11). This indicates the potential for more shared regulatory control of genes often in a more cell-specific manner. The other 1,694 novel microRNAs shared 893 seeds, not present on any human microRNAs reported in miRBase (v21).

We then validated 9 novel microRNAs from 6 different cell types or tissue by PCR (Fig. 3C). As an example of potentially interesting novel microRNAs, JHU_ID_23828-3p was validated by PCR and is located in intron 8 of the *EGFL7* gene, approximately 900 bp from miR-126, and within the same pri-miRNA transcript (Chang et al. 2015) (Fig. 3D,E). Unsurprisingly, due to the specific high abundance of miR-126 in endothelial cells, JHU_ID_23828-3p was also significantly more abundant in four endothelial cell lines (avg. 3,673 reads) than 30 other cell types (avg. 8 reads; p=0.001, Mann-Whitney U test, Fig. 3F). Additionally, while miR-126 is highly conserved throughout chordates, JHU_ID_23828 is only conserved among primates (Fig. 3G).

### Wide variation in the distribution of isomiRs

Due to the inexact cutting of Dicer and Drosha and nucleotide additions/modifications, a collection of mostly similar sequences (with most diversity on the 3’ end) make up the isomiR family of a microRNA (Morin et al. 2008; Neilsen et al. 2012). IsomiR families can be comprised of hundreds of different sequences, but most sequences that constitute an isomiR family are templated length variants of the canonical (consensus) sequence and additional nucleotides added to the 3’ end. We evaluated the isomiR distributions of 126 primary cell and 82 cancer/immortalized cell samples.

We evaluated both 3’ length variants ±4 bases from the reported canonical sequence (miRBase v21) and variations in the 5’ nucleotide starting location, which would affect the predicted seed sequence of the microRNA (Fig. 4A, Supplemental Table S12). The most abundant isomiR was widely variable between cells and often incongruent with the expected sequence. In primary cells, 556 microRNAs had reads sufficient for analysis. A comparison of reads, all with the same 5’ starting location, revealed the most abundant isomiR to always be the canonical sequence for only 182 (33%) microRNAs. There were 204 (37%) microRNAs in which the miRBase v21 canonical sequence was never the most abundant sequence (Fig. 4A). This includes miR-10a-5p in which a 1 bp shorter sequence was the dominant species in 111 of 112 samples and miR-140-3p in which the dominant species was 2 bp longer in 91 of 113 samples (Supplemental Table S12).

**Figure 4.**
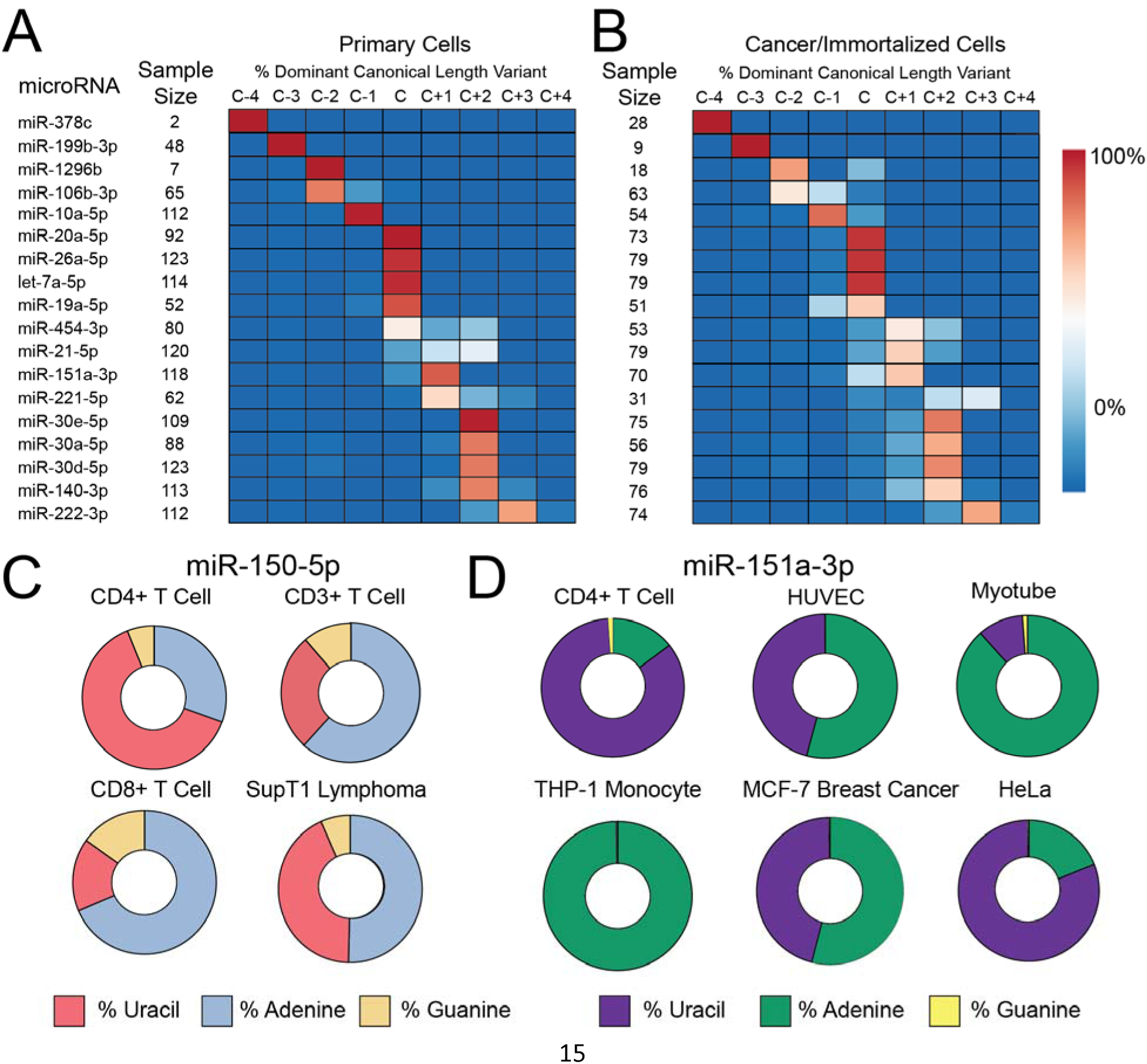
IsomiRs are a challenge to characterizing microRNA levels. (A) Among primary cells, the most abundant (dominant) sequence for many microRNAs differs in length from the canonical “C” miRBase.org v21 sequence by up to 4 bases (C-4 to C+4). Between cell types, length diversity is also present as evidenced by microRNAs that are not entirely of one length. Eighteen representative microRNAs from 556 in total. (B) The general features of microRNA length among cancer/immortalized cells are similar, but the microRNA processing in these cells skews towards randomness. See also Supplemental Figure 10. (C) miR-150-5p, a lymphocyte-specific microRNA, shows a diversity of non-templated nucleotide addition at the +1 site on the 3’ end. Cytosine is the templated (genomic) nucleotide at this position and is not shown. (D) miR-151a-3p, a ubiquitous microRNA, has extreme variation in the first non-templated nucleotide addition. Cytosine is again the templated nucleotide at this position.

Across the primary cells, 84 microRNAs also had more abundant reads for template sequences that started proximal or distal to the canonical 5’ starting position (Supplemental Table S13), which is distinct from the 3’ changes reported in Fig 4A. This included miR-199b-3p (+1 shift), miR-181c-3p (+1 shift) and miR-302a-5p (+3 shift), all of which had highly abundant reads containing a completely different seed sequence than the one currently assigned with strong implications for the targeting of genes (Tan et al. 2014). Although technical factors may be responsible for some variation between cell types, the data clearly demonstrates a need to revise our understanding of the appropriate canonical microRNA sequences for better reproducibility and computational target prediction(Mestdagh et al. 2014; Agarwal et al. 2015).

We then ascertained the nucleotide identity of the non-templated 3’ addition at the +1 position from the most abundant canonical isomiR reported above. Across the 126 cell types, 56% of non-template additions were adenines, followed by 41% uracils, 1% guanidines and 3% cytosines (Neilsen et al. 2012). Between cell types, these values were highly variable, with additional non-templated adenines ranging from 28% (iPSC neurons) to 82% (H9 differentiated cells) of all additional nucleotides. As this may reflect sequencing or batch effect, we investigated just the 32 different primary cells that we generated in a shared batch and observed a two-fold range from 31% (dermal neonatal fibroblast) to 69% (bronchial epithelium) suggesting a real biological phenomenon (Supplemental Fig. S8).

Finally, we assessed what percentage of all reads from a given microRNA family were assigned to the most abundant sequence. On average, only 45% of all microRNA reads for each isomiR group were assigned to the most abundant isomiR sequence (Supplemental Fig. S9). This was quite variable and rarely >90% suggesting methods that fail to acquire isomiRs significantly underestimate the presence of microRNAs in a variable fashion (Mestdagh et al. 2014).

We then investigated the isomiRs of 82 samples from 35 different cancer (or immortalized) cell types. Here we found subtle differences in the canonical microRNAs relative to primary cells. Only 22% of microRNAs (versus 33% of primary cells) had the most abundant isomiR as the miRBase v21 canonical sequence in 100% of cells. The diversity of most abundant microRNA sequences between samples (as characterized by Shannon entropy) among cancer/immortalized cells was a ∼20% increase in disorder over primary cells (p<0.005 Wilcoxon rank sum test) suggesting increased Dicer and Drosha miscleavages. (Fig. 4B, Supplemental Fig. S10, Supplemental Table S14). Nontemplated adenosines (62%) and uracils (30%) were again the dominant 3’ modifications in immortalized/cancer cell samples, but could vary widely between both primary and cancer/immortalized cells as a reflection of either biological differences or technical factors (Fig. 4C,D).

## Discussion

Here we provide the first comprehensive delineation of cell-specific expression patterns of human microRNAs. These ubiquitous and cell-specific patterns of microRNA expression, identified by RNA-seq data, are further supported by matching super-enhancer data and highlight that there are many fewer ubiquitous microRNAs than currently believed. Of key importance is the specific expression patterns of certain microRNAs. Namely, miR-451a and miR-144 are exclusively expressed in red blood cells, yet, because blood is found in all tissues, they have been inappropriately assigned a variety of functions in epithelial and mesenchymal cells based on misinterpreted tissue-level data (Kent et al. 2014; Halushka 2016).

Having this encyclopedic knowledge of microRNA localization provides additional benefits. We provided one example, using adipose, of how cell-specific patterns can deconvolute complex tissue expression patterns. As more data is added to this growing cellular microRNAome, we can effectively work to reduce expression heterogeneity in tissue samples across large studies (McCall et al. 2016). This will improve the interpretation of tissue microRNA expression levels, which, to date, has significantly muddied our understanding of microRNA localization and biologically-relevant function (Kent et al. 2014).

This study also has implications for the measurement and manipulation of microRNA expression. By using cell, not tissue data, we could observe cell-specific differences in the isomiR composition of microRNAs. For 205 common microRNAs, the most abundant sequence did not match the reported sequence in mirBase.org. This difference, which has previously been reported on a smaller scale (Morin et al. 2008), can have an important effect on PCR and hybridization based strategies that may target a secondarily abundant microRNA in the isomiR family, altering the reported expression level of a microRNA. This could explain some of the variability of microRNA expression across methods demonstrated by the miRQC project (particularly between RNA-seq and hybridization approaches) (Mestdagh et al. 2014) and impact the biologic activity of mimics and inhibitors relative to the true microRNA 5p end (Guo et al. 2014).

There are important limitations to this work. Because much of the data is taken from public sources, in which the RNA-seq has been performed across different platforms and with different sequencing methods, significant technical variation in microRNA expression is present. Despite our robust normalization methods, and requirement of >1 million microRNA reads, technical factors certainly drive some of the variation and clustering in these samples. As well, some of these cell types have few to no replicates, and it will be important to continue adding data for these cells. It is also unknown the extent to which microRNA expression patterns of cultured cells match cells *in vivo*. We strongly emphasize that the novel microRNAs described are putative as we have strong concerns about the methodology of current novel microRNA prediction programs and believe they have high false positive rates. New methods, taking into account our collective knowledge of known microRNA structures and isomiR families, must be considered for the next generation of novel microRNA detection algorithms (Backes et al. 2016). Finally, we are only moving towards a complete cellular microRNAome as many cell types (hepatocytes, neutrophils, pneumocytes) were not available for this study.

These microRNA expression patterns from 42 cell types are the first step towards a complete understanding of microRNA expression across all cells and establishing a human cell atlas. Our data also demonstrate general consistencies, but not without some concerning differences, between primary cells and immortalized/cancer cell lines. This work brings a new realization to the importance of cellular microRNA localization and enhances our understanding of this powerful regulatory RNA species.

## Methods

### Cell isolation and sequencing methods

Twenty-nine cell types were obtained from Lonza and cultured according to the manufacturers specifications for no more than 6 passages (Supplemental Table S15). Primary coronary endothelial cells, smooth muscle cells and fibroblasts were isolated and cultured in ECM or SMC media (ScienCell) from a 29 year old man and primary aortic endothelial cells were isolated and cultured from a 10 year old girl as described (McCall et al. 2011a). Red blood cells were isolated from whole blood by centrifugation at 900G for 10 minutes at room temperature and then pipetted and collected. Colonic epithelial cells were obtained by flow sorting (BD FACSAria II) of EpCAM+ cells through a modification of the protocol of (Dalerba et al. 2007). T lymphocytes were obtained by flow sorting of a homogenized spleen sample for CD3+ cells. Cortical neurons were grown from iPSCs using the methods described (Xu et al. 2016). RNAs were isolated with the miRNeasy kit (Qiagen) according to the manufacturer’s protocol. RNA integrity was assessed using Agilent BioAnalyzer and the RNA concentrations were measured using a NanoDrop 2000 UV-Vis Spectrophotometer. Small RNA libraries were prepared using the Illumina TruSeq Small RNA Library Preparation kit according to the manufacturer’s protocol or purified using a Pippin Prep with a 3% Agarose Gel Cassette (Sage Science) and a size selection of 122-157bp. Multiplexed sequencing was performed as single read 50 base pair, using rapid run mode, and v2 chemistry on HiSeq2000 or HiSeq2500 systems (Illumina) at either the Genome Technology Center at the NYU School of Medicine or the Next Generation Sequencing Center at the Johns Hopkins University School of Medicine.

### Differentiation of hES cells into dopamine neurons

H1 human embryonic stem cells (Wi Cell, Madison, WI) were cultured using standard protocols on inactivated mouse embryonic fibroblasts. Differentiation of hES cells to dopamine neurons was done as described (Kriks et al. 2011). Single cells hES cells were cultured on matrigel coated plate at a density of 40,000 cells/cm^2^ in SRM media containing growth factors and small molecules including FGF8 (100ng/ml), SHH C25II (100ng/ml), LDN-193189 (100nM), SB431542 (10μM), CHIR99021(3μM) and Purmorphamine(2μM) for the first five days. Over the next six days, cells were maintained in neurobasal medium containing B27 minus vitamin A, N2 supplement along with LDN193189 and CHIR99021. In the final stage, they were made into a single cell suspension and seeded at a density of 400,000/cm^2^ on polyornithine and laminin coated plate in a neurobasal media containing B27 minus Vitamin A, BDNF(20ng/ml), GDNF(20ng/ml), TGFB1(1ng/ml) ascorbic acid(0.2mM), cAMP(0.5mM) and DAPT(10μM) till maturation (approx. 60 days).

### Publically available RNA-seq data

Sequence Read Archive and Array Express were searched for the terms “human” and “microRNA” or “miRNA” and the records were evaluated for any human primary cell type, cancer cell line, transformed/immortalized cell line or normal human tissue. These generally represented the “control” materials in experiments. In total, 655 sra files were downloaded and converted into fastq files using fastq-dump of the SRA Toolkit. Sequencing was performed on Illumina systems (Genome Analyzer IIx, HiSeq2000, HiSeq2500, miSeq) and AB SOLiD Systems. Solexa colorspace data was converted to standard fastq format using SoLiD2Std.pl. Data searches and collection ended on 2/8/16. The cell line H1264, which is a lung carcinoma cell line, has been reported as being cross contaminated with H157, which is a separate human lung carcinoma cell line (ICLAC.org). However, in the context of the way data from this cell line was used, that distinction is of no consequence here.

### microRNA annotation via miRge

miRge was used as described (Baras et al. 2015). Briefly, miRge removes sequence adapters and performs quality control through cutadapt (Chen et al. 2014). Then, reads are collapsed together and undergo a 5 step alignment to customized RNA libraries utilizing Bowtie and designed to optimally capture microRNAs and their isomiRs. For microRNAs with high sequence similarity (ex. hsa-let-7a-5 and hsa-let-7c-5p), miRge reports them together (ex. hsa-let-7a-5p/7c-5p). The generally used command line for miRge was perl miRge.pl –adapter illumina –species human –CPU 8 –SampleFiles a.fastq,b.fastq… In all, 694 RNA-seq fastq files were run in batches or individually. Prior to the run, the presence and type of adapter was noted for each fastq file. A variety of sequencing methods resulted in a range of adapters used. For some fastq files, adapters were removed using the standalone version of cutadapt. A consensus adapter sequence could not be determined for 45 samples and the sequences were trimmed to 21 bp using the cutadapt –u command (ex. $ cutadapt <FILE>.fastq -u -14 -o <FILE>_cut.fastq for a 35 bp read length). These samples were excluded from isomiR analyses. The 159 samples that had less than 1 million microRNA reads were excluded and are not represented in the data. We also removed 5 tissue samples (SRR1635903-8) with extreme technical skewing of microRNA reads (>60% of all reads were microRNA let-7b-5p).

### DEXUS Analysis

The DEXUS algorithm was used to fit a mixture of five negative binomial distributions to the RNA-seq counts from all cell-type samples (dexus R/BioC package version 1.14.0). We then selected miRNAs that had at least one highly expressed distribution (highest mean >50,000). The most likely distribution from which each miRNA / sample value came (called responsibilities in the dexus package) were used as a discretized measure of expression. Distributions with a mean less than 2,500 were merged into a background / unexpressed distribution. This type of expression discretization has been shown to greatly reduce batch effects when combining data across studies and technologies (McCall et al. 2011b; McCall et al. 2014).

### Remove Unwanted Variation (RUV) Normalization of microRNA samples

The Remove Unwanted Variation algorithm(Risso et al. 2014) using replicate samples (RUVSeq R/BioC package version 1.8.0) was used to estimate five latent factors separately for combined primary cell and cancer cell line data and tissue data. Replicate samples were defined using biologically-based clusters of tissues and cells. We verified the ability of the five estimated latent factors to capture and adjust for batch effects by examining biological clusters comprised of multiple experiments. The RUV normalized data clustered by biological cluster and not experimental batch.

### t-Stochastic Neighbor Embedding

t-SNE was performed using the Rtsne package (version 0.11) in R on RPM corrected cell data and RUV normalized data for primary cells, cancer/immortalized and tissue samples with perplexity set at 10 after evaluations of perplexity values of 1-40 for each (van der Maaten and Hinton 2008). All RUV data was normalized to summed counts and log2 transformed.

### Determining the ubiquity of microRNAs

All 162 primary cell runs were collapsed into their 46 unique cell types keeping the maximum RPM value for each microRNA. This method was replicated for the 26 tissue types. The frequency of a microRNA being >100 RPM across each common cell type was determined in a histogram.

### Calculation of 5p, 3p dominance

The sum of each microRNA’s RPM value across all 162 primary cell runs was generated. microRNAs that had fewer than 1,000 summed RPMs were excluded. Equivalent levels represent the reads for 5p and 3p being within 10% of each other.

### Super-enhancer analysis

Super-enhancer genomic location data was obtained from the dbSUPER website (Khan and Zhang 2016) for 11 cells (primary or cancer/immortalized) in which there was matched RNA-seq data. As microRNA RPM data can be variable between samples, the specific samples used were SRR5127214, SRR5127200, SRR1264358, SRR1575597, SRR1200888, SRR5127213, SRR020286, SRR5127233, SRR873410, SRR1055962, and SRR5127217. The distance between 939 microRNA loci (hg19) and all superenhancers was determined and only those of distance <40kb to a microRNA loci were evaluated. The RPM (log2) of the mature microRNA strand was obtained for each genomic microRNA loci. Because some microRNAs are expressed from multiple genomic locations (ex. let-7a-1, let-7a-2 and let-7a-3 on chromosomes 9, 11, and 22), and there is no way to distinguish the genomic source of the mature microRNA, we assign the mature miRNA expression value (let-7a-5p) to all sites. We caution that the activity of these super-enhancers on adjacent genes and microRNAs is generally unknown and these reported correlations are not proof of activity of the super-enhancer on the microRNA.

### CIBERSORT analysis

The CIBERSORT (Newman et al. 2015) web application: cibersort.stanford.edu was used to create a signature gene matrix using the following parameters: a maximum condition number of 20, qvalue threshold of 0.5, and between 5 and 50 signature genes per cell-type. This signature gene matrix was then used to estimate the composition of tissue samples from the METSIM study (Civelek et al. 2013).

### Immortalized/Cancer vs Normal Cells

RUV corrected microRNA data was plotted using the pheatmap function in R. A H7 ESC sample was used as an outgroup for each correlation. A MAplot was generated for the average of 12 primary fibroblast cell cultures vs. 3 immortalized fibroblast cell cultures. A separate MAplot was generated for 8 primary T cell cultures vs. 14 T cell leukemias/lymphomas.

### Novel microRNA discovery

All reads of length 18-25bp that were initially unmapped were collected using a python script from each appropriate run (479 files). Samples that were trimmed to 21 bp in the initial miRge run were excluded. The SRA was searched for all instances of Argonaut (Ago) precipitated RNA, identifying 105 reads from 19 tissues and cells (Supplemental Table S16). This includes 43 reads used in (Londin et al. 2015). All reads of length 18-25 bp were also taken from the ∼1.7 billion unmapped Ago Clip-Seq reads. Of note, distinguishing linker sequences in this dataset was not necessarily feasible, and 54% of reads were adjusted by cutadapt to only 21 or 22 bp lengths using the command described above, likely resulting in an over identification of Ago-bound microRNAs. Both of the 479 standard small microRNA RNA-seq unmapped samples and the 108 Ago reads were processed in miRDeep2 for novel microRNA detection and aligned to the human genome (GRCh38/hg38). microRNA locations were compared to known repeat elements using the RepeatMasker track from the UCSC Genome Browser. A subset of the 21,338 initial microRNA loci and all 2,724 microRNA loci were evaluated using NovoMiRank and comparing the calculated z-scores to the z-score values of all miRBase versions (Backes et al. 2016). The 2,724 microRNA loci of novel and ‘orthologous’ miRNAs were compared to the 3,707 novel microRNAs described by (Londin et al. 2015) and all Ago-related discovered microRNAs. Any overlap between genomic locations indicated a shared loci. Summary statistics were generated for the dataset. pri-miRNA localization was performed using the UCSC tracks generated by Chang TC *et al* (Chang et al. 2015).

### Comments on novel microRNA localization

For many novel putative microRNAs, the sequence reads overlapped, with several bases of extension/difference between samples. Thus the exact chromosome location of each novel microRNA is from a single sample of the collection and may not reflect the best absolute location on GRCh38/hg38. The microRNAs designated as .5p/.3p were named as such as each of these microRNA loci was identified in more than one sample, with two different pre-miRNA structures designating the sequence to the 5p or 3p arm. There were no “passenger” reads to distinguish the correct structure, so either location remains plausible.

### Novel microRNA seed region analysis

The seed region (bases 2-7) were identified on each purported novel microRNA. Due to some 5’ ambiguity between samples, the seed bases could differ for the same microRNA found between two different samples accounting for 3,697 seeds of the dominant strand from the 2,724 novel microRNA loci.

### Amplification of novel microRNAs

Amplification of novel PCRs was based on the stem-loop method of reverse transcriptase (RT) followed by PCR amplification of the microRNA as performed (Londin et al. 2015). miR-21-5p, a ubiquitous and abundant microRNA, was used as a positive control for all RNA sources. All RT primers, PCR primers, and PCR conditions are provided in Supplemental Table S17.

### IsomiR analysis

A perl script was generated that took the mapped.csv file and counted the reads for each microRNA’s canonical microRNA sequence (from miRBase v21), length variants from -4 to +4 bp around the canonical sequence, additional canonical sequences, and the number of reads for non-templated (non-genomic) nucleotide additions (A,G,C,U) to the maximal count canonical length variant. Only microRNAs with 1,000+ total reads and >10% of reads that were canonical length variants were evaluated. 126 primary cell samples (15,897 microRNA reads) and 82 cancer cell samples from 35 unique cancer (or immortalized) cell types had appropriate data for analysis. The nomenclature C-4, C-3, C-2, C-1, C, C+1, C+2, C+3, C+4 indicate the length of the dominant templated microRNA species relative to the canonical (C) sequence.

### Entropy analysis

The distribution of the lengths of miRNA species detected from a given miRNA family (locus) was characterized relative to the length of the canonical sequence as (≤-4, -3, -2, -1, 0, 1, 2, 3, ≥4). The degree of disorder in the cell line culture samples was characterized relative to the median Shannon entropy of the primary cell culture samples, in which this was used as the reference point in the calculation of percent maximal information loss (relative and normalized entropy calculation).

## Data Access

Novel RNA-seq data generated in this project is available through Bioproject PRJNA358331 and samples SRR5127200-36 & SRR5139121 in SRA.

## Acknowledgements

The authors thank C. Porter and L. Blosser for their work in isolating colonic epithelium, Gourav Dey for illustrations, Srikanth Manda for cell culture and bioinformatics, both Josh Hertel and Tai C. Huang for RNA isolation, and Adriana Heguy and the NYUMC Genome Technology Center. MKH was supported by the American Heart Association [13GRNT16420015]. The NYUMC Genome Technology Center is partially supported by the Cancer Center Support Grant, P30CA016087, at the Laura and Isaac Perlmutter Cancer Center. AP was supported by NCI’s Clinical Proteomic Tumor Analysis Consortium initiative (U24CA160036 and U24CA210985). PL was supported by the National Fund for Scientific and Technological Development, FONDECYT 1151008, Government of Chile. This work was supported by grants from MSCRFII-0429 and MSCRFII-0125 to V.L.D. 2013-MSCRF-0054 to J-C.X., 2014-MSCRF-0665 to M.K, and NIH/NINDS NS67525, NS37388 to T.M.D and V.L.D. T.M.D. is the Leonard and Madlyn Abramson Professor in Neurodegenerative Diseases.

## Author Contributions

M.K.H., A.P. and K.H.B. conceived the project. M.S.K., P.L., T.M.D., J.X., M.K., V.L.D. generated cell data. A.H.P., C.J.M., A.S.B., Y.L., D.E.A., A.Z.R. and M.N.M. performed computational analysis. M.A. validated novel microRNAs, M.K.H. and M.N.M. wrote the paper.

